## Supplementary figures and images for "Insect Detect: An open-source DIY camera trap for automated insect monitoring"

### S1 Fig

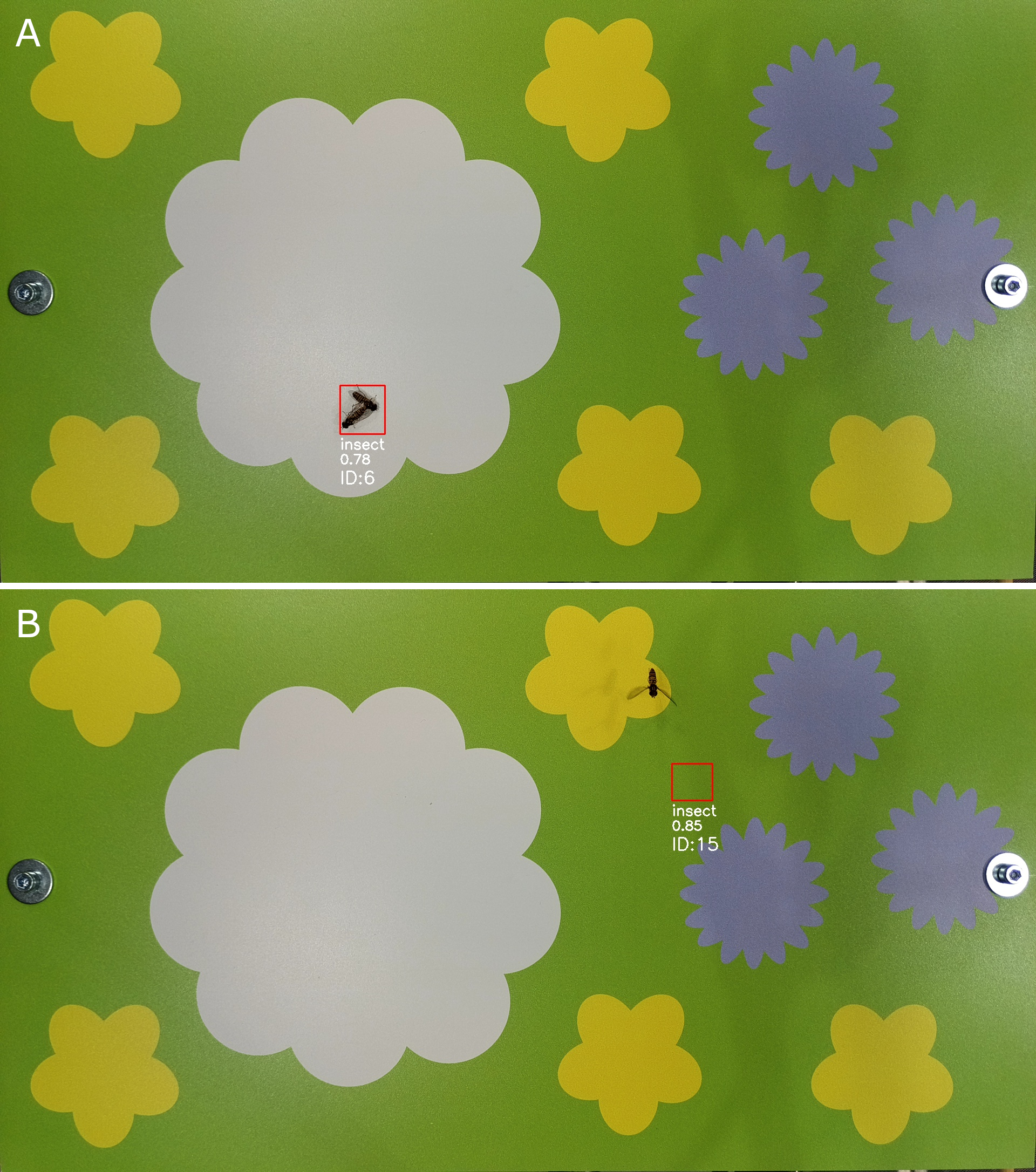

### S2 Fig

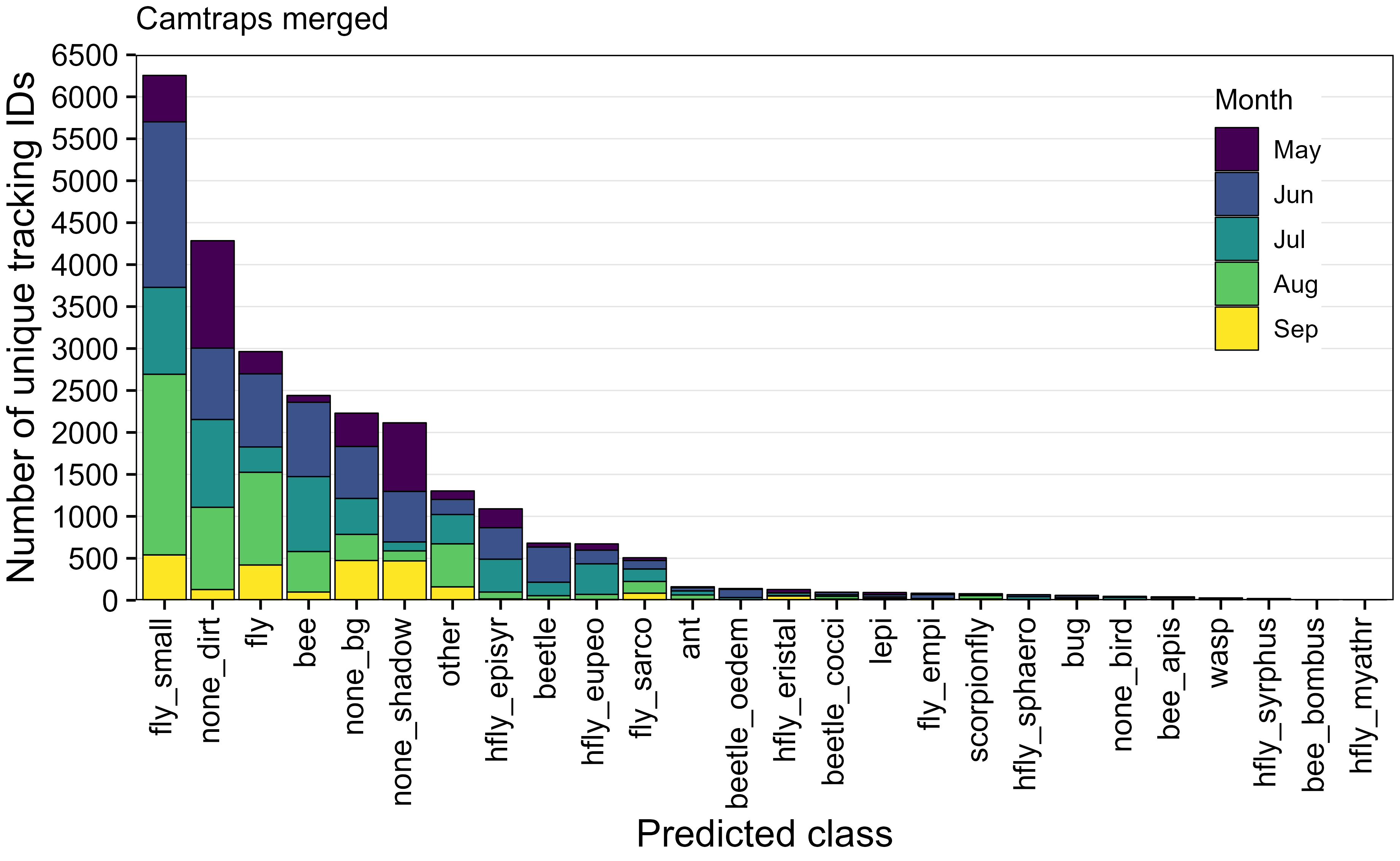

### S3 Fig

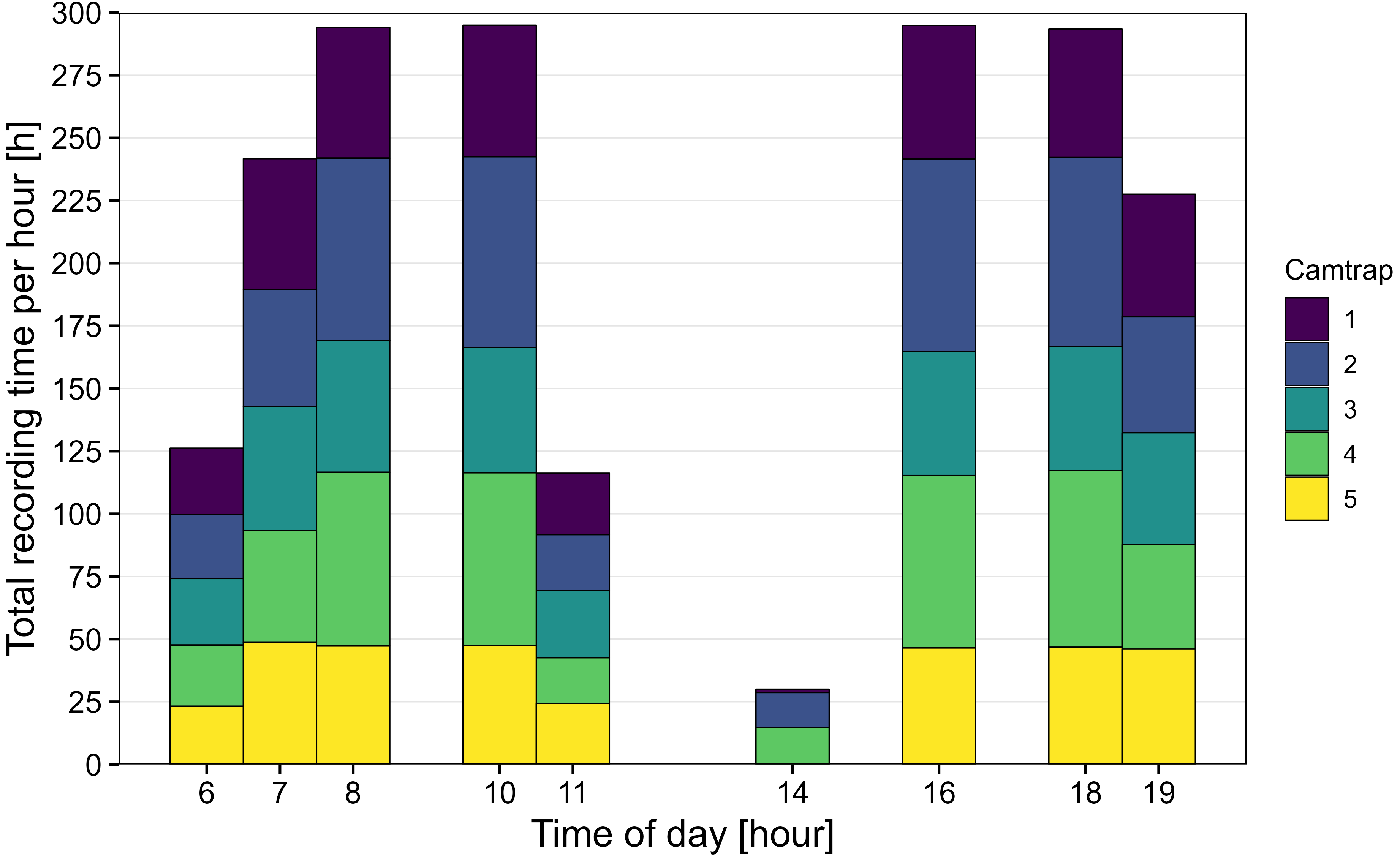

### S4 Fig

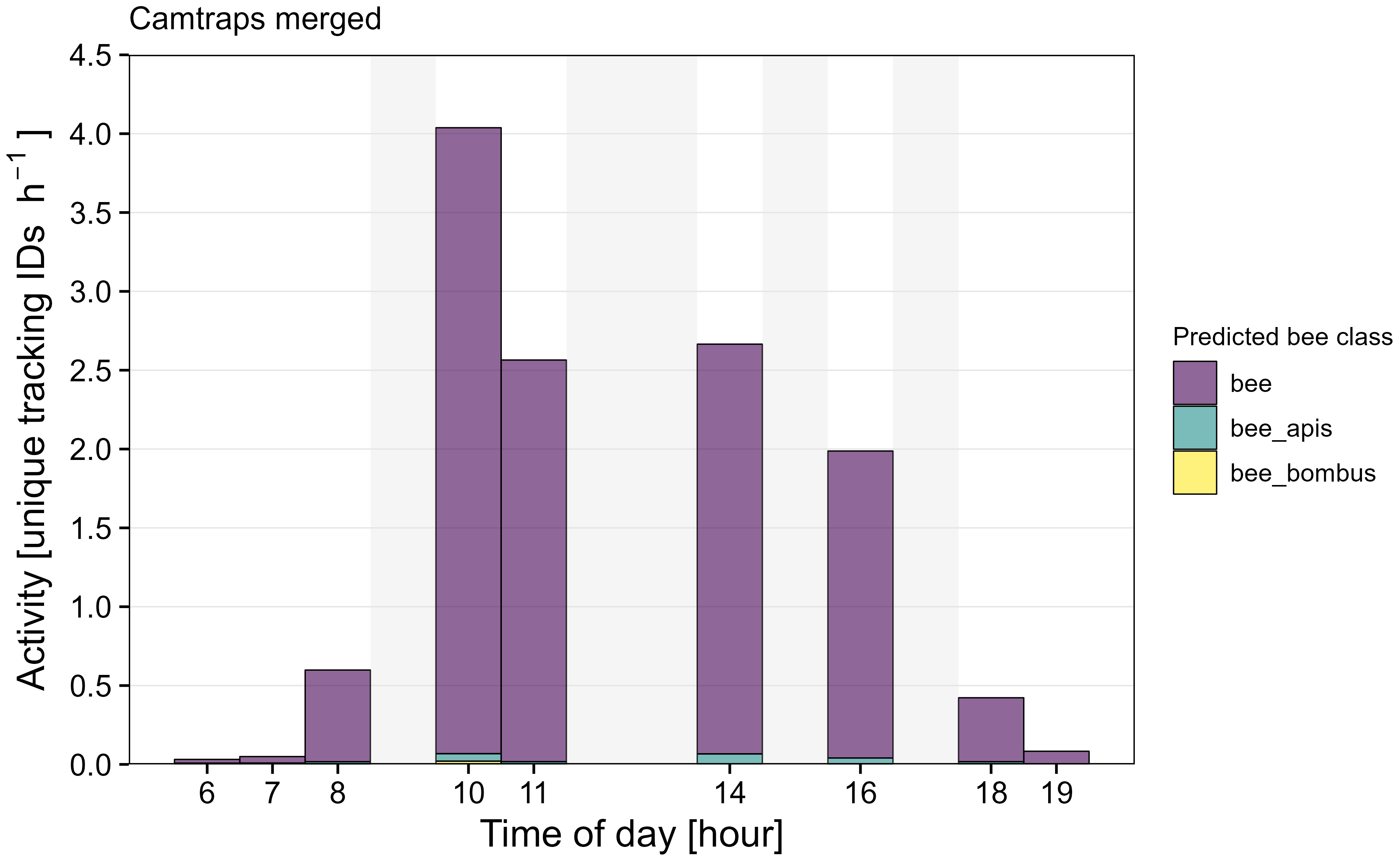

### S5 Fig

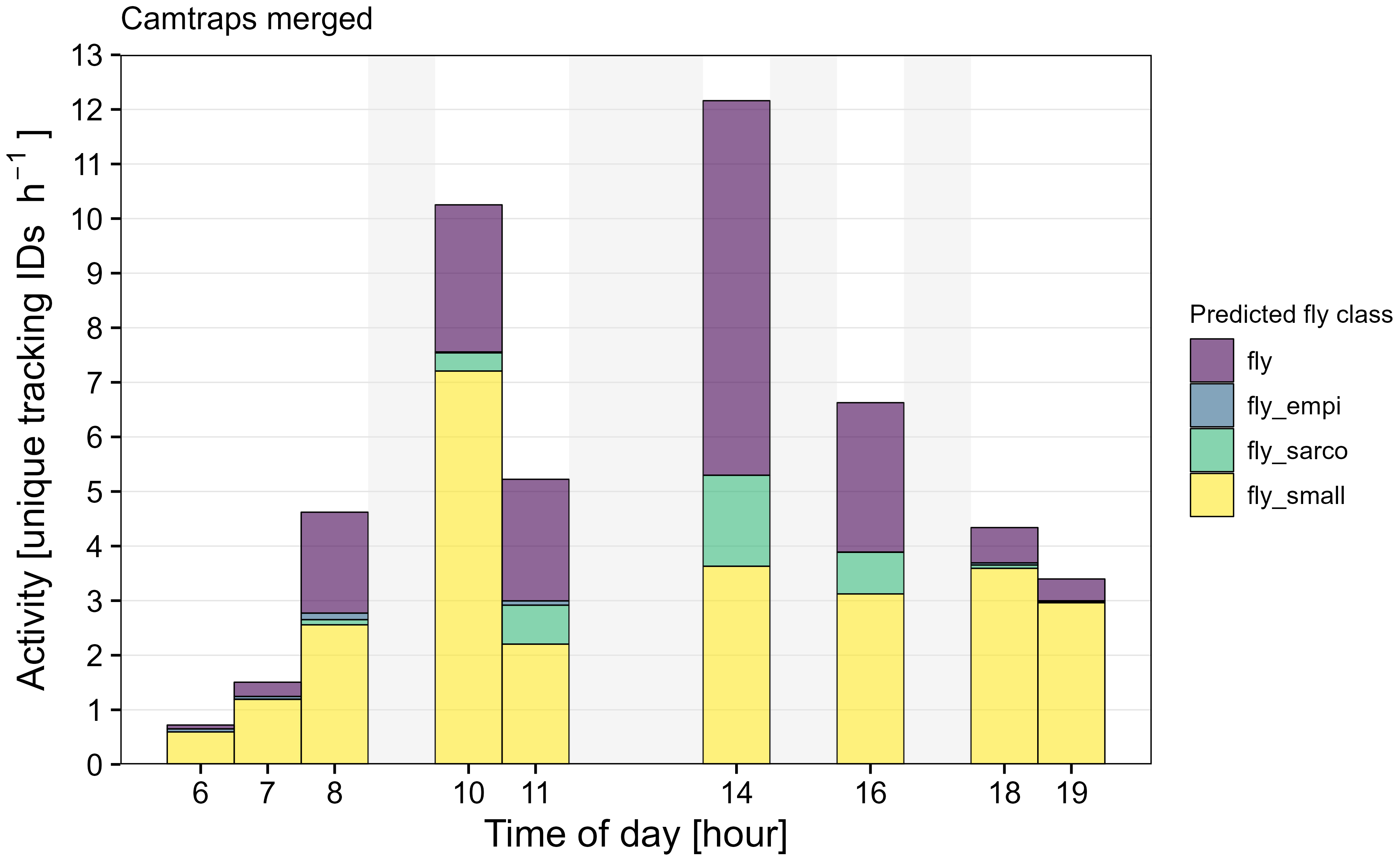
