## Supplementary material for "Insect Detect: An open-source DIY camera trap for automated insect monitoring": S1 Table

**S1 Table. Description of the 27 classes from the image dataset that was used to train the insect classification model.**

| **Class name** | **Description** |
| --- | --- |
| ant | Formicidae (Ants) |
| bee | Anthophila (Bees), excluding *Apis mellifera* and *Bombus* sp. |
| bee_apis | *Apis mellifera* (Western honey bee) |
| bee_bombus | *Bombus* sp. (Bumblebees) |
| beetle | Coleoptera (Beetles), excluding Coccinellidae and some visually distinct Oedemeridae |
| beetle_cocci | Coccinellidae (Ladybugs) |
| beetle_oedem | Oedemeridae, only visually distinct species, mostly *Oedemera* sp. |
| bug | Heteroptera (True bugs), excluding *Graphosoma italicum* |
| bug_grapho | *Graphosoma italicum* (Italian striped bug) |
| fly | Brachycera excluding Empididae, Sarcophagidae, Syrphidae and small Brachycera |
| fly_empi | Empididae (Dagger flies) |
| fly_sarco | Sarcophagidae (Flesh flies) |
| fly_small | small Brachycera, visually distinct to bigger Brachycera |
| hfly_episyr | Hoverfly species *Episyrphus balteatus* |
| hfly_eristal | Hoverfly species *Eristalis* sp., mainly *Eristalis tenax* |
| hfly_eupeo | Hoverfly species *Eupeodes corollae* and *Scaeva pyrastri* (could include other, visually similar hoverfly species) |
| hfly_myathr | Hoverfly species *Myathropa florea* |
| hfly_sphaero | Hoverfly species *Sphaerophoria* sp., mainly *Sphaerophoria scripta* |
| hfly_syrphus | Hoverfly species *Syrphus* sp. (could include other, visually similar hoverfly species) |
| lepi | Lepidoptera (Butterflies) |
| none_bg | Images with no insect: background (platform) |
| none_bird | Images with no insect: bird sitting on platform |
| none_dirt | Images with no insect: leaves and other plant material, bird droppings |
| none_shadow | Images with no insect: shadows of insects or surrounding plants |
| other | other Arthropods, including various Hymenoptera, Symphyta, Diptera, Orthoptera, Auchenorrhyncha, Neuroptera, Araneae |
| scorpionfly | Scorpionfly species *Panorpa* sp. |
| wasp | Vespidae, mainly *Vespula* sp. and *Polistes dominula* |

The images were sorted to the respective class by considering taxonomic and visual distinctions. In some cases, clear taxonomical separations are difficult from images only and the decision to sort an image to the respective class was based more on visual distinction.
