## Supplementary material for "Insect Detect: An open-source DIY camera trap for automated insect monitoring": S2 Table

**S2 Table. Comparison of different classification model architectures and hyperparameter settings supported by YOLOv5 classification model training.**

| **Model** | **Image size**  **[pixel]** | **Batch**  **size** | **Number**  **epochs** | **Top-1**  **accuracy^val^** | **Top-1**  **accuracy^test^** | **Precision^val^** | **Precision^test^** | **Recall^val^** | **Recall^test^** | **F1 score^val^** | **F1 score^test^** |
| --- | --- | --- | --- | --- | --- | --- | --- | --- | --- | --- | --- |
| EfficientNet-B0 | 128 | 64 | 5 | 0.977 | 0.969 | 0.975 | 0.965 | 0.976 | 0.962 | 0.975 | 0.963 |
| EfficientNet-B0 | 128 | 64 | 10 | 0.98 | 0.977 | 0.981 | 0.977 | 0.974 | 0.974 | 0.977 | 0.975 |
| EfficientNet-B0 | 128 | 64 | 15 | 0.979 | 0.972 | 0.98 | 0.971 | 0.973 | 0.964 | 0.976 | 0.967 |
| EfficientNet-B0 | 128 | 64 | 20 | 0.98 | 0.972 | 0.979 | 0.971 | 0.974 | 0.967 | 0.976 | 0.969 |
| EfficientNet-B0 | 128 | 64 | 25 | 0.976 | 0.97 | 0.975 | 0.966 | 0.97 | 0.962 | 0.972 | 0.963 |
| EfficientNet-B0 | 128 | 32 | 15 | 0.977 | 0.973 | 0.971 | 0.974 | 0.97 | 0.963 | 0.97 | 0.968 |
| EfficientNet-B0 | 128 | 96 | 15 | 0.981 | 0.973 | 0.982 | 0.973 | 0.976 | 0.969 | 0.979 | 0.971 |
| EfficientNet-B0 | 128 | 128 | 15 | 0.981 | 0.972 | 0.982 | 0.97 | 0.979 | 0.968 | 0.98 | 0.969 |
| EfficientNet-B0 | 128 | 96 | 20 | 0.981 | 0.974 | 0.981 | 0.975 | 0.975 | 0.97 | 0.978 | 0.972 |
| EfficientNet-B0 | 128 | 128 | 20 | 0.976 | 0.971 | 0.978 | 0.97 | 0.97 | 0.966 | 0.974 | 0.968 |
| EfficientNet-B0 | 96 | 64 | 15 | 0.977 | 0.969 | 0.974 | 0.969 | 0.971 | 0.961 | 0.972 | 0.965 |
| EfficientNet-B0 | 160 | 64 | 15 | 0.98 | 0.978 | 0.98 | 0.976 | 0.97 | 0.974 | 0.975 | 0.975 |
| YOLOv5s-cls | 128 | 64 | 10 | 0.966 | 0.953 | 0.965 | 0.955 | 0.953 | 0.939 | 0.958 | 0.946 |
| YOLOv5s-cls | 128 | 64 | 15 | 0.965 | 0.954 | 0.963 | 0.954 | 0.947 | 0.943 | 0.954 | 0.948 |
| YOLOv5s-cls | 128 | 64 | 20 | 0.965 | 0.95 | 0.965 | 0.95 | 0.949 | 0.935 | 0.956 | 0.942 |
| ResNet-50 | 128 | 64 | 10 | 0.963 | 0.954 | 0.963 | 0.951 | 0.949 | 0.941 | 0.956 | 0.945 |
| ResNet-50 | 128 | 64 | 15 | 0.961 | 0.963 | 0.958 | 0.96 | 0.946 | 0.954 | 0.951 | 0.956 |
| ResNet-50 | 128 | 64 | 20 | 0.963 | 0.954 | 0.963 | 0.955 | 0.949 | 0.942 | 0.955 | 0.947 |

All models were trained on a custom dataset with 21,000 images (14,686 in train split) and default hyperparameters. Metrics are shown on the dataset validation split (4,189 images) and dataset test split (2,125 images) for the converted models in ONNX format.
