## Supplementary material for "Insect Detect: An open-source DIY camera trap for automated insect monitoring": S3 Table

**S3 Table. Metrics of the EfficientNet-B0 insect classification model, validated on a real-world dataset.**

| **Class** | **Images** | **Top-1 accuracy** | **Precision** | **Recall** | **F1 score** |
| --- | --- | --- | --- | --- | --- |
| all | 97671 | 0.827 | 0.712 | 0.817 | 0.698 |
| ant | 54 | 0.796 | 0.088 | 0.796 | 0.159 |
| bee | 658 | 0.907 | 0.712 | 0.907 | 0.798 |
| bee_apis | 292 | 0.884 | 0.902 | 0.884 | 0.893 |
| bee_bombus | 18 | 1.0 | 0.783 | 1.0 | 0.878 |
| beetle | 972 | 0.197 | 0.575 | 0.197 | 0.293 |
| beetle_cocci | 27 | 0.667 | 0.75 | 0.667 | 0.706 |
| beetle_oedem | 89 | 0.831 | 0.574 | 0.831 | 0.679 |
| bug | 383 | 0.441 | 0.621 | 0.441 | 0.516 |
| bug_grapho | 1 | 1.0 | 1.0 | 1.0 | 1.0 |
| fly | 10660 | 0.927 | 0.754 | 0.927 | 0.832 |
| fly_empi | 1 | 1.0 | 0.004 | 1.0 | 0.008 |
| fly_sarco | 2500 | 0.939 | 0.988 | 0.939 | 0.963 |
| fly_small | 36478 | 0.902 | 0.988 | 0.902 | 0.943 |
| hfly_episyr | 682 | 0.963 | 0.966 | 0.963 | 0.965 |
| hfly_eristal | 568 | 0.979 | 0.938 | 0.979 | 0.958 |
| hfly_eupeo | 392 | 0.962 | 0.917 | 0.962 | 0.939 |
| hfly_myathr | 39 | 0.897 | 1.0 | 0.897 | 0.946 |
| hfly_sphaero | 157 | 0.981 | 0.846 | 0.981 | 0.909 |
| hfly_syrphus | 38 | 0.868 | 0.825 | 0.868 | 0.846 |
| lepi | 21 | 0.762 | 0.485 | 0.762 | 0.593 |
| none_bg | 1454 | 0.872 | 0.172 | 0.872 | 0.287 |
| none_bird | 13 | 1.0 | 0.022 | 1.0 | 0.044 |
| none_dirt | 17529 | 0.733 | 0.912 | 0.733 | 0.813 |
| none_shadow | 18376 | 0.788 | 0.831 | 0.788 | 0.809 |
| other | 3433 | 0.386 | 0.576 | 0.386 | 0.462 |
| scorpionfly | 2767 | 0.896 | 0.999 | 0.896 | 0.945 |
| wasp | 69 | 0.493 | 1.0 | 0.493 | 0.66 |

The model was trained on a custom dataset with 21,000 images (14,686 in train split), scaled to 128x128 pixel, to 20 epochs with batch size 64 and default hyperparameters. Metrics are shown on a real-world dataset (97,671 images) for the converted model in ONNX format. All images were classified and subsequently verified and sorted to the correct in class in the case of a wrong classification by the model. A dummy image was added for each of the classes “bug_grapho” and “fly_empi”, as no images of both were captured during the selected recording period. Results for both classes must be ignored.
